# Functional connectome harmonics capture early brain organization and maturity in neonates

**DOI:** 10.64898/2026.03.25.714150

**Authors:** Aylin Rosberg, Isabella L. C. Mariani Wigley, Aaron Barron, Ilkka Suuronen, Harri Merisaari, Elmo P. Pulli, Silja Luotonen, Ru Li, Wajiha Bano, Ashmeet Jolly, Hilyatushalihah Kholis Audah, Niloofar Hashempour, Elena Vartiainen, Hasse Karlsson, Linnea Karlsson, Richard A. I. Bethlehem, Jakob Seidlitz, Jakub Vohryzek, Joana Cabral, Dafnis Batalle, Morten L. Kringelbach, Selen Atasoy, Jetro J. Tuulari

## Abstract

The functional organization of the human brain is established early, yet the ontogeny of its large-scale functional gradients remains unclear. Using resting-state fMRI data from 714 neonates in the Developing Human Connectome Project, we mapped neonatal brain gradients via functional connectome harmonics (FCH). We identified adult-like sensory-to-multimodal and cognitive gradient patterns present at birth. Applying three FCH-derived metrics—entropy, power, and energy—we found that power and energy were higher in term-born compared to preterm neonates, while entropy was elevated in preterms. These metrics predicted up to ~30% of postmenstrual age, indicating their biological relevance. Our findings reveal that the neonatal brain possesses a robust gradient architecture underpinning early functional organization, offering novel biomarkers for assessing brain maturity and the impact of prematurity on neurodevelopment.

## Introduction

The macroscale organization of the human brain is increasingly understood not as a collection of discrete regions, but as a series of spatially continuous gradients. These gradients represent a hierarchical axis that transitions from primary sensory-motor regions to higher-order associative cortex, providing a structural framework that supports complex cognitive functions. Recent evidence demonstrates that brain networks can be reconstructed from structural and functional gradients, and that gradient patterns are an intrinsic property of human brain organization across multiple scales ^1–5^.

Despite the fundamental importance of this hierarchical architecture in adults, the ontogeny of these gradients remains largely unexplored. It is unclear whether gradient patterns are an emergent property of postnatal experience or a pre-configured template present at birth. Here, we characterize functional gradients of the neonatal brain using functional connectome harmonics (FCH) ^6^. Using a large, high-quality dataset of 714 term- and preterm-born neonates from the Developmental Human Connectome Project (dHCP, postmenstrual age (PMA) at scan 26–45 weeks), we demonstrated that a near-adult-like functional topology is already established at the earliest stages of life, comparable to that observed in a subsample of adults from the Human Connectome Project (HCP, N = 99). We further show that these neonatal gradients align with both functional connectivity and meta-analytic functional activation patterns ^7^, and can be used to predict established brain network patterns.

While the theoretical basis and research of brain gradients have matured considerably ^8^, there remains a translational gap: the field lacks standardized, derived measures that can be applied to clinical cohorts. Here, we build on previously introduced gradient-based metrics designed for applied neuroimaging, entropy, power, and energy ^9–13^, and validate these measures by demonstrating, for the first time, their sensitivity as neural biomarkers that can accurately predict PMA and identify signatures of neurodevelopmental vulnerability in infants born preterm. Together, our findings suggest that the gradient-based architecture of the brain is a robust, early-emerging feature of human neurodevelopment that offers a novel and quantitative approach for assessing brain maturity and pathology.

## Results

### Functional connectome harmonics reveal novel gradients in the neonatal brain

The neonatal brain exhibits remarkable levels of anatomical and functional organization, even though it has achieved only approximately 30% of its final volume and its white matter is only partially myelinated ^14,15^. By term-equivalent age, well-known resting state networks such as sensorimotor, auditory, and visual networks are clearly apparent in a nearly mature configuration in both term and preterm-born infants ^16–22^. These networks exhibit little change during subsequent development, whereas higher-order networks encompassing frontal and parietal brain areas are fragmented in neonates and attain their adult-like configuration during the first 24 months after birth ^20,23^. To determine whether a similar level of network maturation applies to gradient patterns, we computed functional harmonics of the group-average connectivity matrix, estimated through parcellation with the AAL atlas that included 90 cortical and subcortical regions. Functional connectivity was quantified as the temporal correlation between the fMRI time series of the cortical and subcortical brain regions. The group-level connectivity matrix was transformed into an adjacency matrix, from which functional connectome harmonics were estimated by eigendecomposition of the corresponding graph Laplacian (Fig.1).

**Figure 1.**
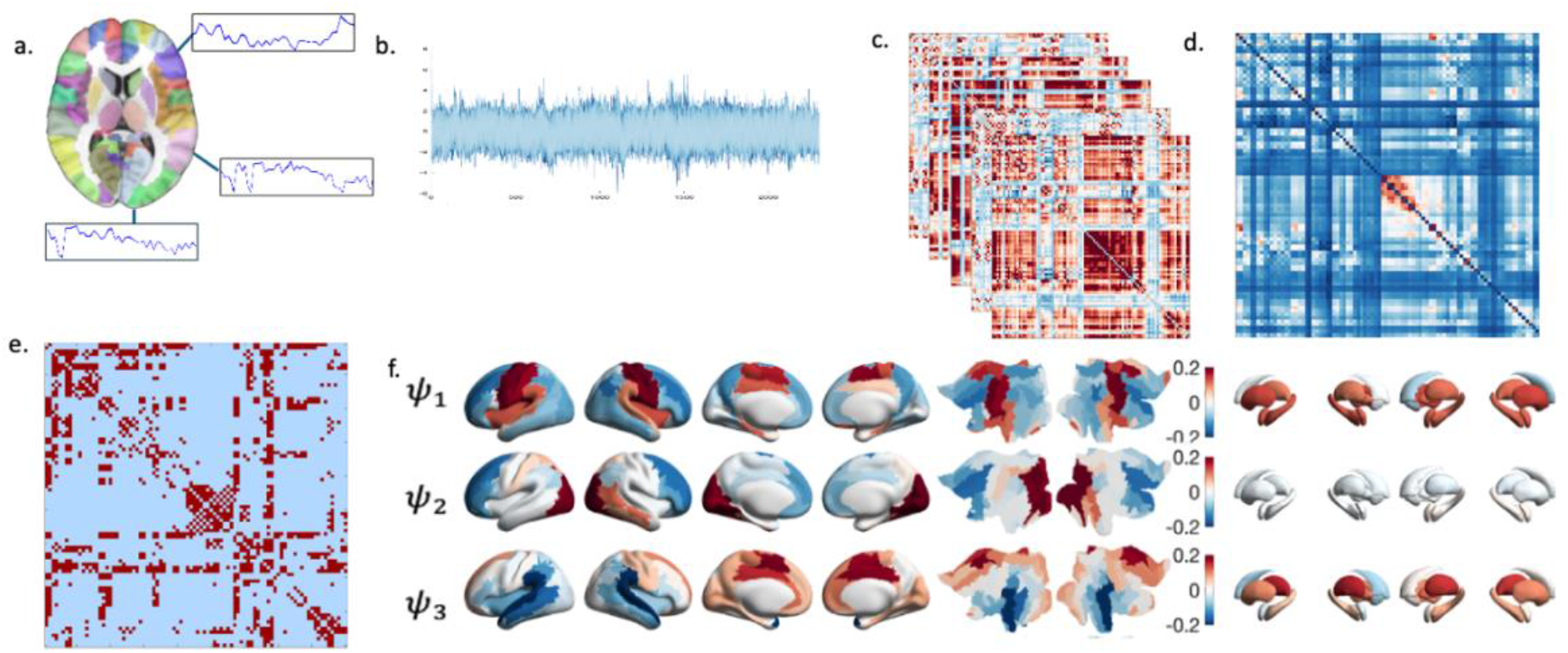
Summary of the workflow: a) individual preprocessed fMRI data (2300 volumes / time points) were parcellated according to the AAL atlas into 90 cortical and subcortical brain regions; b) yielding 90 x 2300 timeseries; c) the time series were z-standardized and used to estimate individual 90 x 90 functional connectome matrices (N = 714); d) a group average connectome matrix was estimated from the individual matrices. e) The symmetrized and binarized adjacency matrices were constructed from the group average connectome matrix using a 10-nearest neighbour (k=10) procedure. f) The functional connectome harmonics were derived as the Laplacian eigenvectors of this adjacency matrix via symmetric Laplacian eigenmaps, (three first FCH are visualized but harmonics were generated across 89 spatial frequencies).

FCH represent spatial patterns across parcellated brain regions, with the sign and magnitude of harmonic values reflecting differences in functional connectivity profiles along each gradient. Based on their correspondence with established cortical gradients, we selected the following neonatal brain harmonics for subsequent analyses: *ψ*_1_ (sensory–multimodal), *ψ*_2_ (anterior–posterior), *ψ*_3_ (medial–lateral), *ψ*_4_ (fronto–temporal), *ψ*_5_ (fronto–parietal), and *ψ*_6_ (interhemispheric) connectome harmonics(Fig.2).

**Figure 2.**
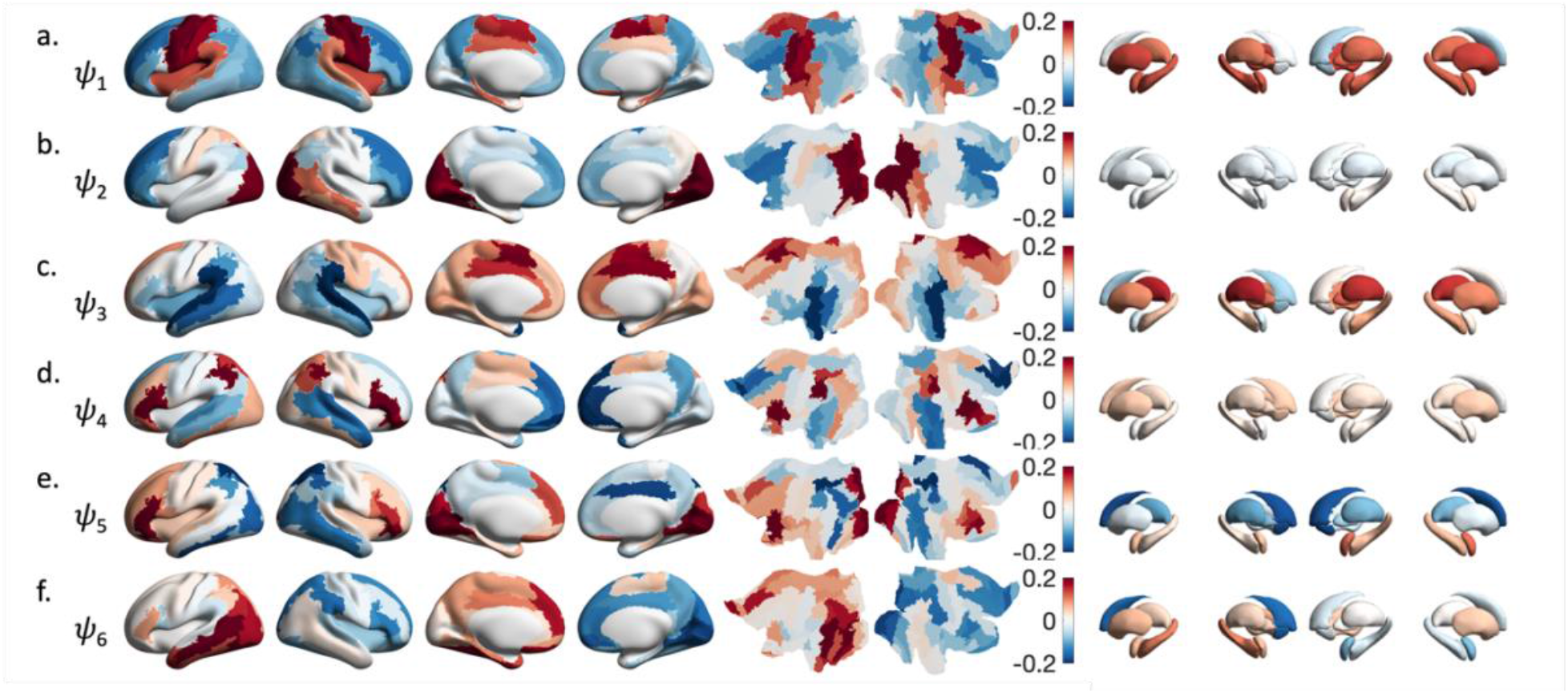
The Functional Connectome Harmonics (FCH) describe the principal gradient patterns of the neonate brain. The first six functional connectome harmonics with each harmonic represented with a letter from a to f; (and numerals *ψ*_1_, *ψ*_2_, …, *ψ*_6_) derived from 714 infants’ fMRI data series averaged into a group mean functional connectivity matrix plotted on the cortical surface and subcortical structures. Left to right: left and right lateral; left and right medial cortical surface plots; unfolded cortical representations of left hemisphere and right hemisphere side by side; and medial and lateral views of subcortical nuclei. By definition, the first functional harmonic with eigenvalue 0 is constant over all subjects and does not contain any spatial information (not shown).

### FCH of the neonate brain associate with adult FCH and known brain gradients

Adult FCH calculated with the same method, showed significant correlations (p<0.05) with neonate FCH, except for *ψ*_5_ (Supplementary Table 1). The correlation analysis between the six neonate FCH and other known cortical gradients revealed that the neonatal functional connectome harmonics are generally well aligned with the organization of adult brain gradients including functional gradients #1–#4 by Margulies et al. ^4^, and first principal component of NeuroSynth (PC1 NeuroSynth) by Yarkoni et al ^24^ (Supplementary Table 2). We found that the *ψ*_1_ sensory-multimodal harmonic showed significant correlations with functional gradients #1 (r = −0.506, p = 0.012) and #2 (r = −0.506, p = 0.015) and PC1 NeuroSynth (r = 0.493, p = 0.012). *ψ*_2_ anterior-posterior connectome harmonic was negatively correlated with functional gradient #1 (r = −0.412, p = 0.027) and positively correlated with functional gradient #2 (r = 0.624, p = 0.012). *ψ*_4_ was linked to functional gradients #3 (r = 0.304, p = 0.012) and #4 (r= 0.388, p = 0.015). *ψ*_5_ was linked to functional gradient #4 (r= 0.368, p = 0.015), and *ψ*_6_ was linked to functional gradient #3 (r= −0.192, p = 0.012). These Pearson correlations were controlled for multiple comparisons across all correlations using the Benjamini–Hochberg false discovery rate (FDR) procedure.

### FCH can predict patterns of dynamic phase locking states

We compared the spatial patterns of the FCH with 6 states of phase-locking captured from the same fMRI data with Leading Eigenvector Dynamics Analysis (LEiDA) using partial least squares regression (PLSR) (Fig. 3). Across all six LEiDA brain states, the first PLSR component based on FCH matrix already accounted for a substantially higher proportion of variance (R^2^ = 59.3–98.6%) than the randomized counterpart (R^2^ = 43.3–54.3%) (Supplementary Table 3). Adding a second component led to near-complete prediction for all LEiDA states with the original harmonics (R^2^ ≥ 99.9%), whereas the randomized harmonics remained markedly less predictive (R^2^ = 61.3–71.9%). These findings indicate that functional connectome harmonics provide a specific and informative basis for predicting dynamic brain states, beyond what can be achieved with a randomized harmonic basis.

**Figure 3.**
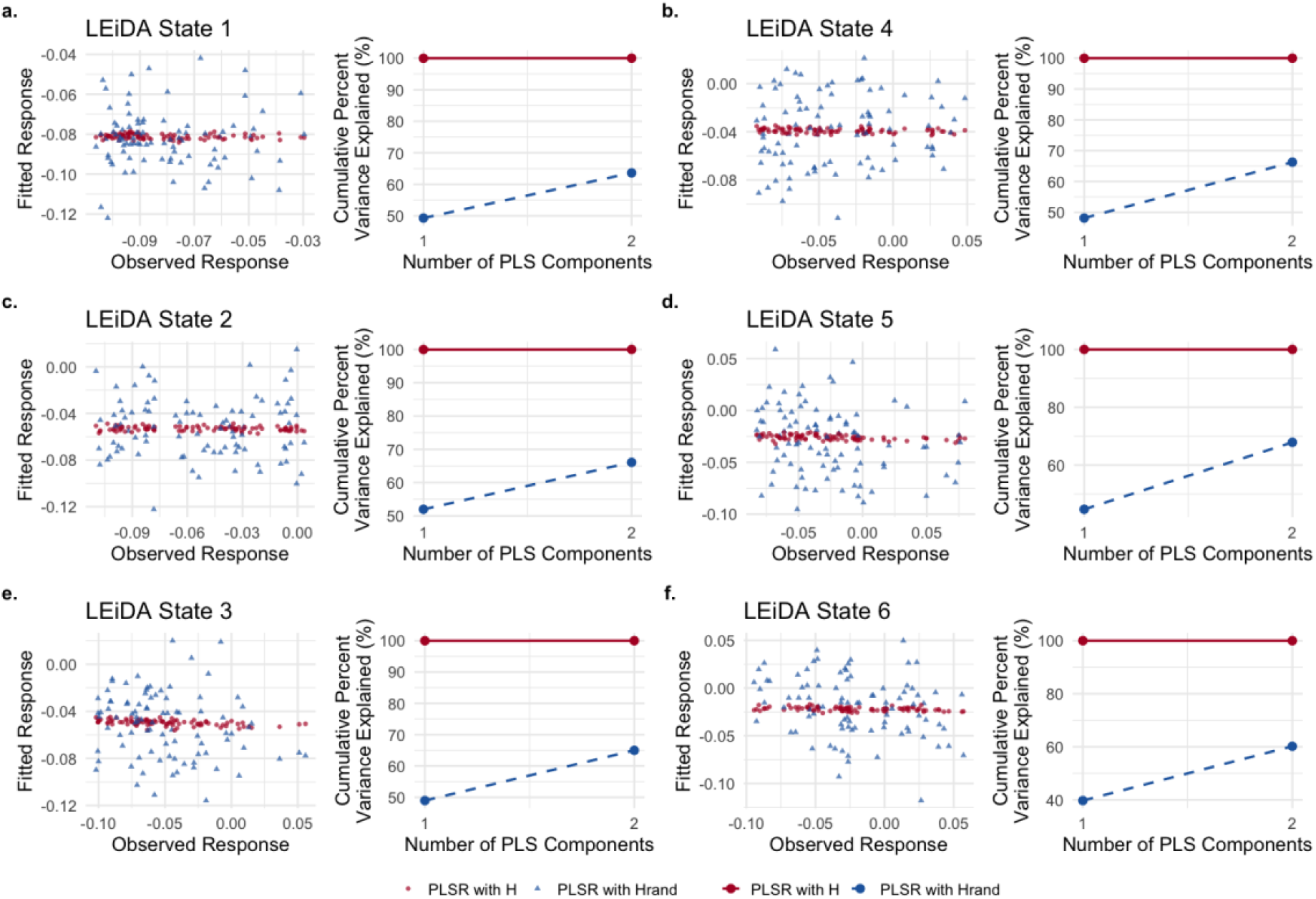
Comparison of model performance across six LEiDA states (each LEiDA state represented with a letter from a to f). (Left Column) Observed vs. Predicted responses for each LEiDA state, showing the predicted responses from PLSR models using the original functional connectome harmonic matrix (H) and its randomized counterpart (Hrand). (Right Column) Cumulative percentage variance explained as a function of the number of PLS components for each LEiDA state. The original harmonic matrix outperformed the randomized version across all states. To assess whether H produced more homogeneous prediction residuals than randomized counterpart, we employed Levene’s test on residuals from 10-fold cross-validation for each brain state. Using two PLS components, H produced significantly lower residual variance compared to randomized harmonics in 5 out of 6 brain states (Levene’s F-test, p < 0.05). Across these significant states, the variance ratio of H_rand/H ranged from 1.63 to 2.65 (mean ± SD: 1.90 ± 0.47).

### FCH-derived brain metrics associate strongly with preterm birth and postmenstrual age

After confirming that the FCH capture meaningful information on brain organization, we proceeded to validate the derived brain metrics. At each time point, we projected the instantaneous ROI activity vector onto the FCH-derived gradient patterns by computing their dot product, yielding a time-resolved measure of gradient expression. These time-resolved coefficients were then summarized across time to derive entropy, power, and energy measures. Entropy quantifies the temporal variability of harmonic projection coefficients, capturing the complexity and degree of disorder in gradient expression over time ^12^. Power is defined as the mean absolute amplitude of harmonic projections across time, reflecting the average strength of expression of each harmonic in brain activity patterns. Energy is defined as the frequency-weighted contribution of each harmonic mode to cortical dynamics, combining its instantaneous power with its intrinsic (eigenvalue-dependent) energy. Previous studies have applied these measures across all (binned) functional connectome harmonics to characterize changes in mental states ^9–13^. Here, we focus specifically on the most prominent harmonics, analogous to deriving principal functional networks from functional connectivity and restricting the modeling to these dominant components.

Entropy, power, and energy were calculated for all neonates in the sample (N = 714). Mann-Whitney U tests were performed to explore whether the metrics differed based on preterm status or sex (Supplementary Table 4). Distinct patterns emerged in the relationships between the derived brain metrics and prematurity. Preterm neonates had lower power and energy compared to term-born neonates, indicating a reduced expression of dominant functional connectome harmonics in the preterm group (Fig.4). In contrast, entropy was higher in preterm-born neonates, indicating more variable and less structured large-scale neural dynamics in the preterm group (Fig.4). While power and energy revealed subtle sex differences only in the first harmonic, entropy did not show any sex differences, highlighting its independence from sex-specific factors (Supplementary Table 4).

**Figure 4.**
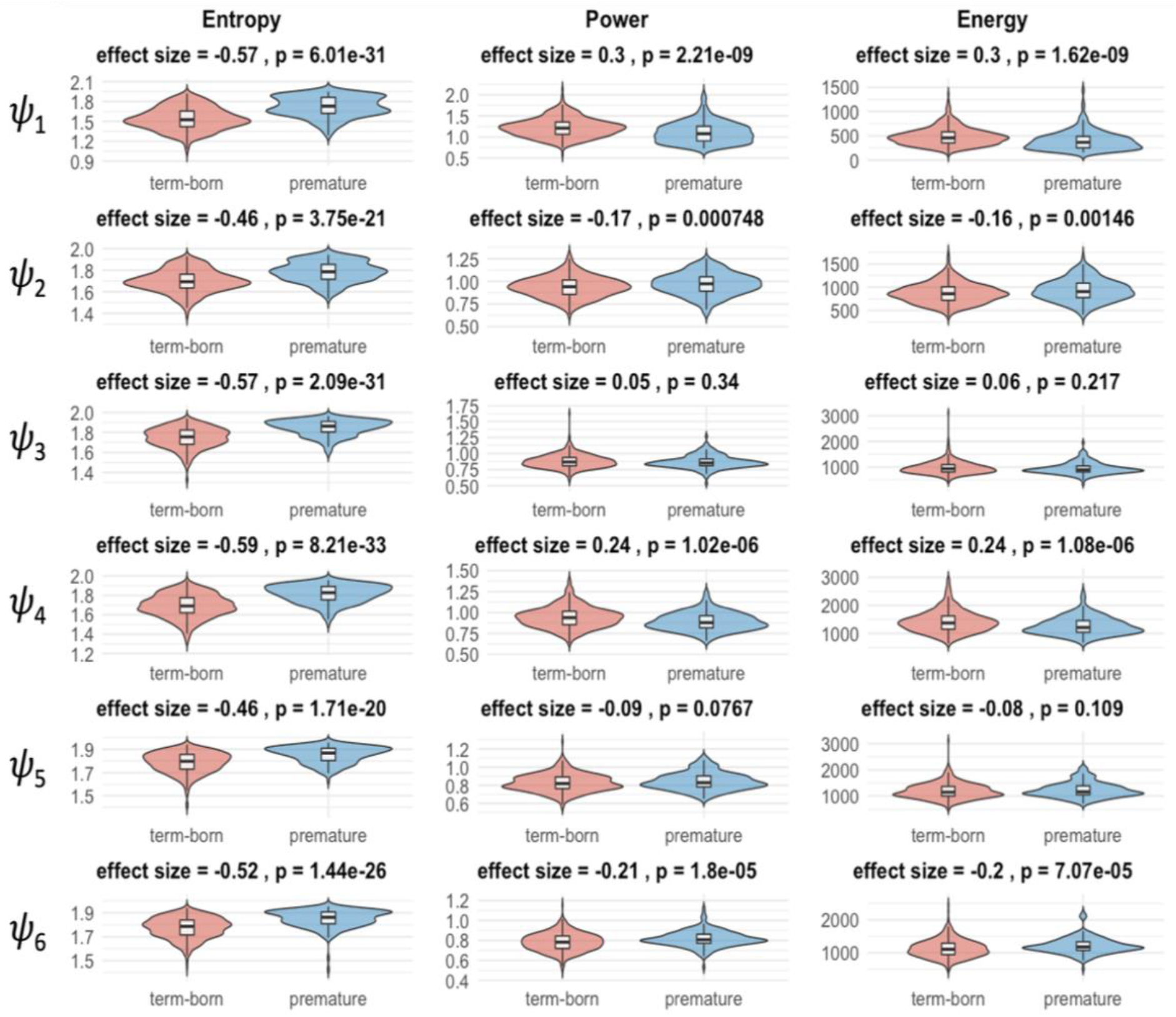
The differences between term-born and preterm-born participants and associations to postmenstrual age at scan for derived measures of the first six functional connectome harmonics (FCH; ψ_1_– ψ_6_, top to bottom). FCH projections on the standardized timeseries (1 x 2300) were used to obtain derived brain measures for entropy, power and energy across time (single scalar value for each participant for each harmonic) all of which are used as derived brain measures in testing the differences between being term-born and preterm-born on brain functional organization. For each harmonic, panels present a violin plot illustrating the distribution of the derived measure (entropy, power, or energy) across participants, grouped by birth status (term-born in red, preterm in blue). The effect size (Rank-biserial correlation coefficient) and the FDR-corrected p-value above each pair of plots are from a Mann-Whitney U test comparing term-born and preterm-born groups.

PMA is tightly associated with brain growth and functional maturation. We used the FCH-derived brain metrics to predict PMA in the whole sample and using only the term-born neonates. Elastic net regression results revealed that power and energy had identical predictive power for PMA (R^2^ = 0.32) and age from birth (AfB) (R^2^ = 0.13), notably higher than that of entropy (PMA R^2^ = 0.24; AfB R^2^ = 0.04). All three metrics predicted PMA much better than they did AfB (Fig.5). Together, these results demonstrate that the FCH-derived metrics are more strongly associated with neurodevelopmental maturation (PMA) than with chronological age since birth (AfB).

**Figure 5.**
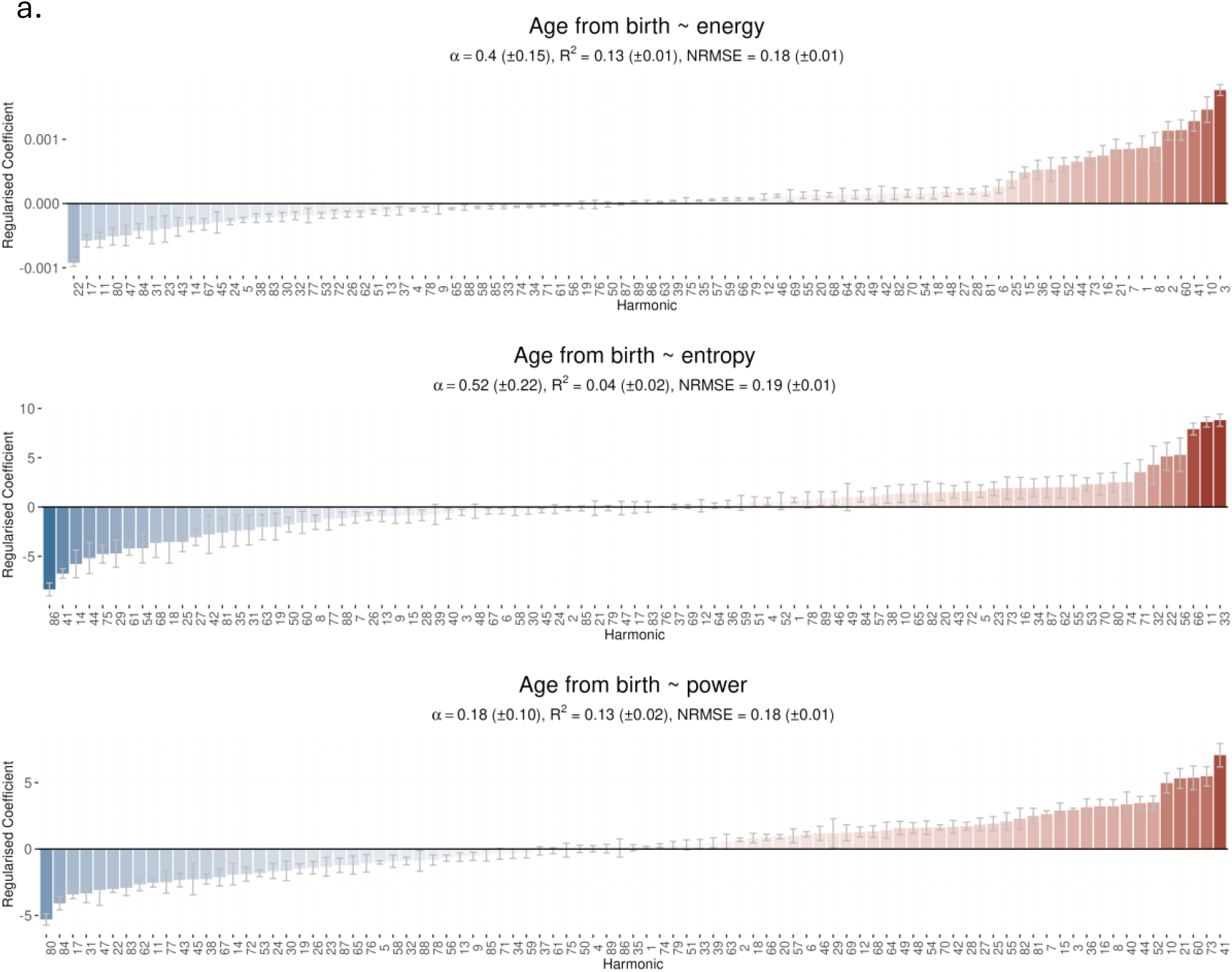
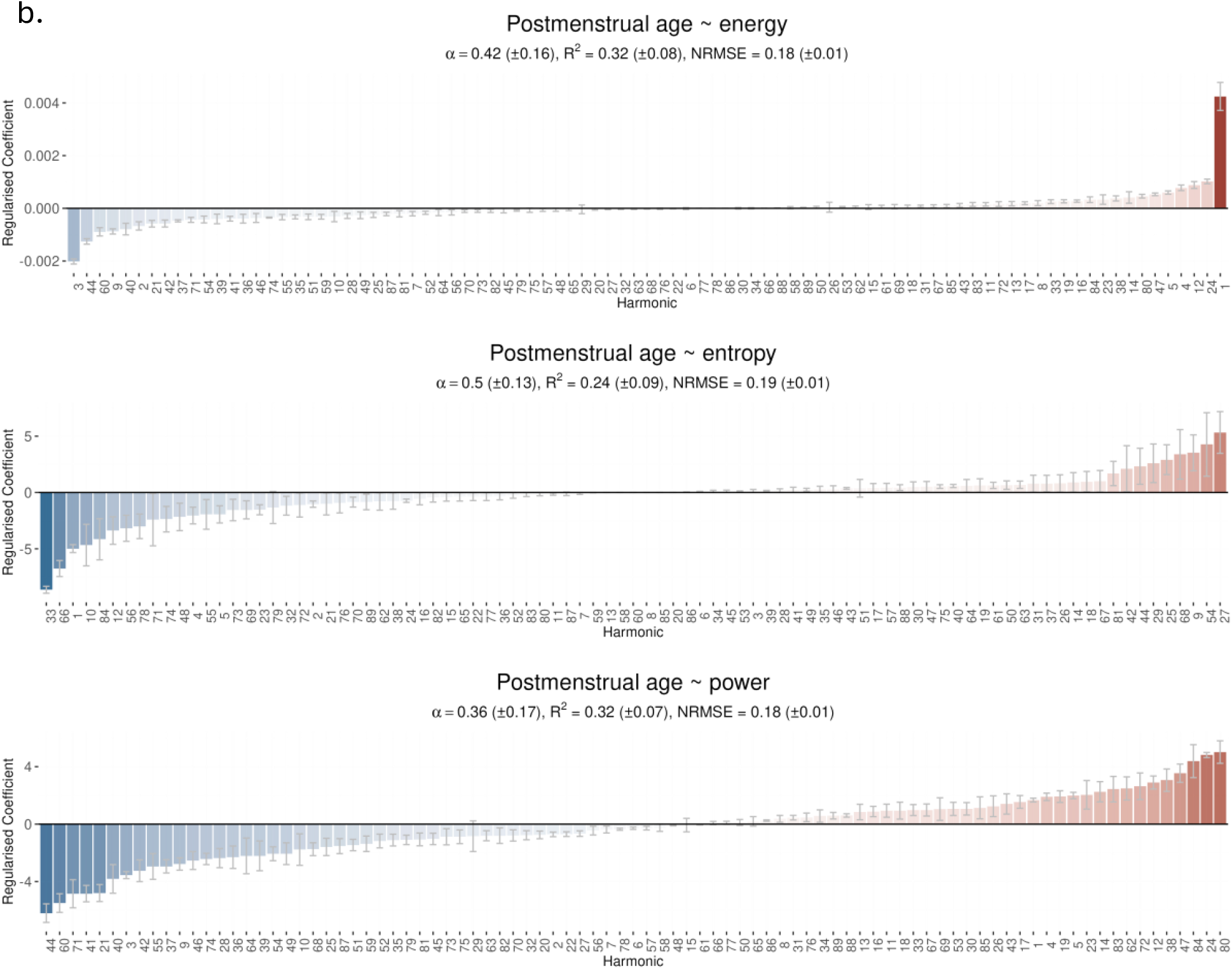
Predictions of age from birth (a.) and postmenstrual age (b.) at scan across all functional connectome harmonics calculated with elastic net regression using three derived measures entropy, energy and power. Bar plots displaying the standardized regression coefficients for each FCH, sorted in ascending order. Positive coefficients (red bars) indicate harmonics with direct associations to the outcome, while negative coefficients (blue bars) indicate inverse relationships.

Further, the predictive performance differed between the full sample and the term-born subgroup across all derived brain metrics. For entropy, prediction of PMA was stronger in the full sample (R^2^ = 0.26) than in term-born neonates alone (R^2^ = 0.15). However, a contrasting pattern was observed for AfB, with increased predictive power in the term-born subgroup (R^2^ = 0.13) compared to whole sample (R^2^ = 0.04). Power showed the highest predictive performance for PMA in the whole sample (R^2^ = 0.33), while prediction was weaker when using only term-born neonates (R^2^ = 0.15). For AfB, the predictive power in term-born subgroup was higher (R^2^ = 0.21). Energy exhibited a pattern comparable to power, with stronger prediction of PMA in the whole sample (R^2^ = 0.32) than in term-born neonates and of AfB in the term-born subgroup (R^2^ = 0.21). Overall, the results suggest that different brain metrics capture different temporal aspects of early development, with sensitivity shifting from maturational age (PMA) in mixed cohorts to postnatal age (AfB) within the term-born subgroup.

## Discussion

Using functional connectome harmonics, we demonstrate that large-scale gradient organization is already present in the neonatal brain and closely aligns with canonical adult functional gradients. Despite ongoing structural immaturity, neonatal harmonics recapitulate sensory– multimodal, anterior–posterior, and transmodal organizational axes and show significant correspondence with established adult gradients. These harmonics provide a compact and biologically grounded basis for predicting dynamic phase-locking states, substantially outperforming randomized harmonic representations and underscoring their specificity for neonatal brain dynamics. Notably, harmonic-derived brain metrics are more strongly associated with postmenstrual age than with age from birth, suggesting that the expression of large-scale functional gradients primarily reflects intrinsic neurodevelopmental maturation rather than postnatal experience.

Our findings provide a comprehensive characterization of the neonatal brain’s functional architecture through the lens of Functional Connectivity Harmonics (FCH). We demonstrate that the macroscale organizational principles previously identified in adults ^2,25^, specifically the hierarchical transition from sensory-motor to multimodal associative regions are already present at birth. The high correlation between neonatal FCH and adult gradients suggests that the human brain exhibits a harmonic scaffold that is already present at birth. By identifying these modes within the widely-used neonatal AAL atlas, we provide a bridge between theoretical neurophysics and standard developmental neuroimaging, ensuring our methodology is accessible for high-throughput analysis on standard computational hardware. A critical insight from this study is the relationship between static harmonic modes and dynamic phase-locking states. By showing that FCH can predict states defined by LEiDA, we establish that the “connectome harmonics” are not merely descriptive spatial patterns but serve as the fundamental basis functions that constrain transient neural dynamics. This suggests that the neonatal brain’s repertoire of functional states is shaped by its underlying harmonic geometry, providing a potential link between structural-functional blueprints and emergent functional organization.

Importantly, we show sensitivity of harmonic-derived metrics—entropy, power, and energy— in predicting postmenstrual age (PMA) and differentiating between preterm and term-born neonates. The ability of FCH metrics to accurately predict PMA points to a largely intrinsic maturational program that proceeds in utero. These metrics thus offer a promising set of neural biomarkers for assessing brain maturation and vulnerability in early life. While the current study focuses on fMRI-based gradients, the spatial properties of these harmonics suggest broad applicability across modalities. Because FCH are anchored to cortical anatomy, they allow for potential integration with EEG and MEG source localization, offering a path toward understanding brain harmonics at much higher temporal resolutions. Future work should investigate how these harmonic “fingerprints” evolve across the first years of life and whether they can serve as early warning signs for neurodevelopmental disorders.

The use of the dHCP data provided us with the largest available neonatal fMRI sample with state-of-the-art sequences, custom head coils, and preprocessing ^26^. This also comes with important limitations, as such settings might be challenging to replicate (for instance, the fMRI data have 2300 time points acquired at a temporal resolution of 0.392 s). We acknowledge that the fMRI images have a low signal-to-noise ratio in inferior cortical regions and across the subcortical regions ^27,28^. We did observe that the overall connectivity from the regions of low SNR to other regions for affected brain areas was comparably low but could also be a normal feature of the neonate brain (although this is hard to confirm due to the paucity of reported connectivity data in prior studies). The functional MRI preprocessing pipeline for dHCP ^26^ accounts for the potential issue of head motion in resting-state fMRI data ^29,30^, and we additionally removed potentially motion corrupted images (and their signal) from the analyses. However, motion also serves as a proxy for the infant’s arousal state, which can influence underlying neural activity ^31–33^ and, consequently, both the modeling and the resulting brain measures.

Studies vary widely in their choice of denoising approaches and exclusion of fMRI volumes with identified motion ^34,35^. For instance, a prior study with the same source population opted for a conservative approach that excluded participants based on motion and used only a subset of the timeseries data with least motion ^33^. Here, the modelling relies on building a connectivity graph through the estimation of group-level functional connectivity matrix. This approach does not necessitate continuous fMRI data, and we thus opted to remove the motion outliers from the data and control for lost temporal degrees of freedom (number of identified motion outliers) per participant in the statistical analyses.

Our investigation establishes FCH as a fundamental organizational principle of the neonatal brain. By demonstrating that the first six FCH mirror the macroscale gradients observed in adults, we show that the brain’s functional hierarchy is a pre-configured blueprint present from the earliest stages of life, rather than a slowly emergent property of childhood.

## Methods

Participants were prospectively enrolled in the dHCP, an observational, cross-sectional Open Science initiative approved by the UK National Research Ethics Authority (14/LO/1169). Written informed consent was obtained from all families prior to imaging. Of note, the enrolment, image acquisition, and preprocessing methods are identical with the prior article, and their description in the methods section has been kept similar for consistency ^33,36^.

### Participants

#### Neonate sample

We included 714 structural-functional datasets acquired from the dHCP neonatal data release 3.0 ^37^. Whenever a participant had multiple scans available, only the first dataset was used. In the study population, there were 519 term-born (282 male, 237 female) neonates who were scanned at 37.4–44.9 weeks PMA and 195 preterm-born (106 male, 89 female) neonates who were scanned at 26.7–45.1 weeks PMA.

### Adult fMRI validation sample

We used the resting-state fMRI data from 100 (45 male, 55 female) healthy young adult participants with mean age 29.5 drawn from the HCP Q3/HCP 900 data release ^38,39^. One participant excluded due to missing data.

### MRI data acquisition

Imaging was performed using a 3T Philips Achieva scanner with modified Release 3.2.2 software, equipped with a specialized neonatal imaging setup comprising a 32-channel phased array head coil and a customized patient handling system (Rapid Biomedical GmbH, Rimpar, Germany). The scanning process and acquisition protocol are optimized for neonates. T2-weighted and inversion recovery T1-weighted multi-slice images were acquired in sagittal and axial stacks with an in-plane resolution of 0.8 × 0.8 mm^2^ and 1.6 mm slices, overlapping by 0.8 mm (0.74 mm for T1-weighted sagittal). For T2-weighted imaging, the parameters were TR/TE = 12000/156 ms with SENSE factors of 2.11 (axial) and 2.60 (sagittal). For T1-weighted imaging, TR/TI/TE = 4795/1740/8.7 ms was used with SENSE factors of 2.27 (axial) and 2.66 (sagittal). Additionally, 3D MPRAGE images were acquired with a resolution of 0.8 mm isotropic and parameters TR/TI/TE = 11/1400/4.6 ms, along with a SENSE factor of 1.2 RL (Right-Left). A high-temporal-resolution fMRI protocol designed for neonates was used, employing multiband (MB) 9 × accelerated echo-planar imaging. Data were collected over 15 minutes with the following parameters: TE/TR = 38/392 ms, 2300 volumes, and a spatial resolution of 2.15 mm isotropic. Detailed information about scanning procedures can be found in the relevant dHCP publication ^37^.

### fMRI data preprocessing

Resting state fMRI data were preprocessed using the dedicated dHCP pipeline optimized for neonatal imaging ^26^. The pipeline includes correction for susceptibility distortions and motion artifacts through slice-to-volume and rigid-body registration ^40–43^. To account for noise contributions from head motion, cardiorespiratory fluctuations, and multiband acquisition, 24 extended motion parameters together with independent noise components identified by ICA and classified using FSL FIX were regressed out ^44^. Denoised data were registered to native T2-weighted space with boundary-based registration ^45^, and subsequently non-linearly warped to the dHCP spatiotemporal template ^46^ using diffeomorphic multimodal registration ^47^.

### Time series extraction and removal of motion outliers

We estimated the motion outliers with fsl_motion_outliers and data points/volumes with DVARS (the root mean square intensity difference between successive volumes) > 1.5 interquartile range (IQR), after motion and distortion correction, were considered as motion outliers, ^26^ in line with prior work ^33^. The number of motion-outlier volumes remaining in the cropped dataset was recorded for each subject and included as a covariate in statistical analyses. The median number of motion-outlier volumes was 38 volumes, range 0–350. We obtained the timeseries of regions of interest as defined in the AAL atlas adapted to the neonatal brain ^48^, which yielded 90 (brain regions) x 2300 (time points) matrices ^36^. We then used a custom MATLAB script to remove the timepoints that were flagged as outliers so that the data that were used for FCH modelling contained only data points that were deemed to be of good quality.

### Modelling the functional connectome harmonics from the fMRI timeseries data

Glomb et al. described a novel method of mapping the functional organization of the adult human brain into a set of cortical FCH ^6^. FCH are estimated by the eigenvectors of the graph Laplacian computed on the graph representation of the functional connectome, and in the original description this was carried out in dense connectomes (59,412 × 59,412 cortical vertices) of the HCP data without any anatomical parcellation. We identified the following challenges to advance the use of FCH in applied studies in developmental neuroscience: 1) The file sizes of gifti/cifti files are large, which could make the computations and data storage challenging; 2) while the use of cifti files is common, it is not mainstream, and the majority of studies use volumetric approaches for estimating the timeseries; and 3) the current use of only cortical data for the FCH modelling is a limitation in terms of many relevant research questions. Finally, direct application of the methodology used in Glomb et al. in the neonatal population is also not straightforward since the neonatal fMRI data have proportionally poorer spatial resolution than adult scans, which means that gifti/cifti files likely contain large amounts of redundant information^49^. To circumvent these challenges, we modified the FCH modelling to accommodate volumetric timeseries of the AAL atlas and subsequent modelling to implement widely adoptable data modelling, reduce computational costs, and improve comparability across other studies as the AAL atlas has been the most frequently used parcellation in neonate studies ^50^.

All data modelling and data visualization were performed using MATLAB version 2024b. We used the group-average functional connectivity matrix as a graph to estimate the eigenfunctions of the graph Laplacian. To achieve this, functional connectivity matrices were generated from the volumetric time series of AAL parcellation for each subject. These individual matrices were then averaged to create a group-level functional connectivity matrix. In constructing the graph representation, 𝒢 = (𝒫, ε) the vertices from AAL parcellation served as nodes, denoted by 𝒫 = {*p*_*i*_| i ∈ 1, …, n}, where 𝒫 represents the AAL atlas parcellation and n = 90 indicates the total number of parcellation regions (nodes) used in this study. The edges ε = {e_ij_|(p_i_, p_j_) ∈ 𝒫 × 𝒫} represented the connections between vertices, which were defined by the correlation values between nodes in the group-average functional connectivity matrix. A binary adjacency matrix *A* = [a_ij_] was created by connecting each node *i* to its *k*-nearest neighbors (with *k* = 10) based on correlation strength. Specifically:

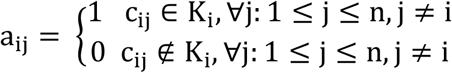

where K_i_ represents the set of the k largest values in row *i* of the group-average functional connectivity matrix. Symmetrization (a_ji_ = 1, *if* a_ij_ = 1) was applied to ensure that **A** is a symmetric, sparse binary matrix which is essential to facilitate eigenvalue decomposition. The diagonal degree matrix **D** was defined such that each entry 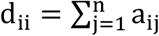 represents the number of connections for node *i*. The graph Laplacian was then computed as **L**_𝒢_ = **D** − **A**, capturing the connectivity structure of the graph and ensuring that rows and columns sum to zero. Connectome harmonics *ψ*_*k*_(*k* = 1, …, *n*) were obtained by solving the eigenvalue problem of the graph Laplacian:

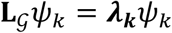

where *λ*_*k*_ is the eigenvalue associated with harmonic *ψ*_*k*_. Harmonics were ordered by increasing eigenvalue, reflecting increasing spatial frequency across the cortical graph.

### Correlations between neonatal FCH, and adult FCH and other adult brain gradients

The relationship between neonatal FCH and adult FCH was investigated by applying the same workflow to 99 adults from HCP data and calculating the correlations between the first six FCH.

The relationships between neonatal FCH and other known cortical gradients were explored by quantifying how similar their values were across corresponding brain regions or vertices. We utilized the Neuromaps toolbox ^7^ to calculate the Pearson’s correlation between the FCH and several well-characterized adult brain maps that are available in Neuromaps including the functional gradient #1– #4 ^51^ which provide low-dimensional axes of resting-state functional connectivity, with the first gradient spanning a principal unimodal–to–transmodal hierarchy from primary sensory–motor to default mode and association cortices, and higher-order gradients (2–4) further differentiating unimodal systems (visual vs somatomotor–auditory) and opposing control/attention networks to default and limbic regions and PC1 NeuroSynth ^24^ capturing large-scale variation in task-evoked activation patterns across cognitive domains.

### FCH predicting LEiDA states

To test whether FCH might be used to predict ‘brain network patterns’, we utilized the LEiDA to model of dynamic brain states. LEiDA is a high temporal resolution method for quantifying brain dynamics and uses phase-locked synchronization of fMRI signal fluctuations in dynamic functional connectivity analysis ^52^. With LEiDA it is possible to detect recurrent BOLD phase-locking (PL) patterns in fMRI signals that closely match the well-known functional networks ^53^. We modeled LEiDA on fMRI signals defined according to the AAL atlas and identified six LEiDA states of the neonate brain (Supplementary Fig.1). The FCH matrix and a randomized set of harmonics were then used to predict the LEiDA states with PLSR. The optimal number of PLSR components was determined using a 10-fold cross-validation. This process was conducted separately for the original functional connectome harmonic matrix (H) and its randomized counterpart (Hrand). The cross-validation criterion minimized the mean squared error (MSE) of prediction to identify the optimal number of components for each model. For all six LEiDA states, the cross-validation results indicated that one to two FCH-based PLS components was sufficient to predict the LEiDA states.

### Statistical analyses testing of differences between derived measures and developmental and demographic variables

The statistical testing and data visualization were carried out in R (versions 4.0.3 and 4.4.0). Entropy was calculated MATLAB (version R2024b) using the approximateEntropy function. Power and energy of connectome harmonics were computed in MATLAB (R2024b). For each subject and connectome harmonic *ψ*_*k*_(*k* = 1, …, *n*), the regional BOLD activity pattern at time *t*, denoted s(*t*), was projected onto the harmonic basis to obtain a projection coefficient:

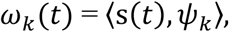

which quantifies the contribution (weight) of harmonic *ψ*_*k*_ to the activity pattern at time *t*.

The instantaneous power of harmonic *ψ*_*k*_ at time *t* was defined as the magnitude of the normalized projection coefficient:

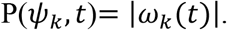

To obtain a single summary measure per subject and harmonic, we averaged the absolute projection coefficients across all time points (T):

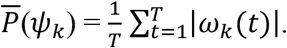

The instantaneous energy of harmonic *ψ*_*k*_ at time *t* was defined as the squared magnitude of the projection coefficient, weighted by the squared eigenvalue *λ*_*k*_ associated with *ψ*_*k*_:

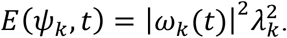

For each subject and harmonic, we then computed the total energy across time points by summing these instantaneous energies over all time points:

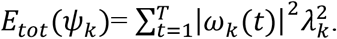

Sex differences and prematurity differences between derived measures were tested with Mann Whitney U test and Rank biseral correlation coeffcient was used as the effect size.

In exploratory analyses, we used five-fold cross validated elastic net regression models to predict PMA and AfB by using entropy, energy and power across all available FCH. Each fold included a 10-fold nested cross validation to choose an optimal α (along a grid from 0 to 1 in intervals of 0.1) and λ (along a grid from 0.01 to 100 in intervals of 0.01) based on the minimal cross validation error. These hyperparameters represent the elastic net mixture between ridge and LASSO regression, and the magnitude of the penalty applied, respectively. Nested cross validation to choose α and λ was performed with the *cv.biglasso()* function from the R package *biglasso*. The elastic net regression model was then fit to the training data with the *biglasso()* function from *biglasso* and used to make predictions in the test data, from which the R^2^ and normalised root mean squared error (NRMSE) values are derived. Further, in exploratory analyses, we employed repeated 10-fold cross-validated Elastic net regression models to predict PMA and AfB using entropy, energy, and power across all 89 available FCH. Elastic net combines L1 (Lasso) and L2 (Ridge) regularization to enable simultaneous feature selection and coefficient shrinkage. Hyperparameter optimization was performed via inner 5-fold cross-validation to select optimal α (l1_ratio; values: 0.1, 0.5, 0.75, 0.9, 0.95, 0.99, 1.0) and λ (alpha; values: 2^-16 to 2^15 on logarithmic scale) based on minimal cross-validation error. The optimized Elastic net models were fit to the training data and used to make predictions on held-out test data, from which R^2^ and mean absolute error values were derived. All features were standardized using z-score normalization prior to model fitting. Analyses were performed separately on the full sample (n=714) and term-born infants only (n=519).

## Code and data availability

The data used in this study were obtained from the Developing Human Connectome Project (dHCP). dHCP data are available through the dHCP data repository: https://nda.nih.gov/edit_collection.html?id=3955

All code used to preprocess the fMRI data have been made available here: https://git.fmrib.ox.ac.uk/seanf/dhcp-neonatal-fmri-pipeline/-/tree/master

## Acknowledgements

The authors wish to acknowledge CSC – IT Center for Science, Finland, for computational resources. Microsoft Copilot was used to improve the grammar of the manuscript that was created by the authors.

The Developing Human Connectome Project was funded by the European Research Council under the European Union Seventh Framework Programme (FP/20072013)/ERC Grant Agreement no. 319456.

